# Mating behaviour, mate choice, and reproduction in *Tyrophagus curvipenis* (Acari: Acaridae)

**DOI:** 10.64898/2026.11.01.763476

**Authors:** Shi-Sen Ye, Keshi Zhang, Zhi-Qiang Zhang

## Abstract

Mating behaviour and mate choice can influence reproductive output, but these aspects remain insufficiently studied in many mite species. This study investigated mating behaviour, mate choice, and the reproductive consequences of repeated mating in *Tyrophagus curvipenis* (Acari: Acaridae). First, we described the mating sequence and quantified the duration of pre-copulation latency and copulation duration. Mating involved initial contact, dorsal mounting, repositioning into a tail-to-tail copulatory position, and sperm transfer. Copulation lasted substantially longer than the pre-copulation latency. Second, we examined mate choice in relation to kinship. Neither males nor females showed a significant preference for siblings or unrelated partners. Third, we examined mate choice in relation to mating status. Males preferred virgin partners, whereas females showed no significant preference between virgin and previously mated partners. Copulation duration was shorter with previously mated partners, although the underlying cause remains unclear. Finally, we examined female reproductive performance under three mating-frequency treatments: one mate, two mates, and unlimited access to mates. Mating frequency did not affect the pre-oviposition period but significantly affected the oviposition period, daily oviposition rate, and total fecundity. Females with unlimited mating opportunities had the longest oviposition period and produced the most eggs. Among females that mated once, longer copulation was positively associated with both oviposition period and total fecundity. These findings characterise mating behaviour in *T. curvipenis* and show that mate choice and mating frequency are associated with different aspects of reproduction. Further work is needed to determine whether these patterns reflect differences in sperm transfer, fertilisation, or other effects of mating.

## 1 Introduction

Mites exhibit extraordinary species richness and evolutionary diversity in morphology, ecology, and reproductive strategies (Norton et al., 1993; Zhang, 2003; Walter and Proctor, 2013b). Their reproductive modes range from asexual reproduction to sexual reproduction and include arrhenotoky, in which males develop from unfertilised eggs (Norton et al., 1993). Among sexually reproducing mites, mating behaviour and sperm-transfer mechanisms also vary considerably (Witte, 1991; Walter and Proctor, 2013a). In Parasitiformes, males often transfer sperm directly into the female reproductive tract using specialised structures associated with the chelicerae (Elbadry and Elbenhawy, 1968; Amano and Chant, 1979; Walter and Proctor, 2013a). Acariformes, on the other hand, exhibit a wider range of mechanisms, including direct insemination using an intromittent organ and indirect sperm transfer through spermatophores, which may be deposited in the presence or absence of a female (Woyke, 1994; Zhang, 1999; Walter and Proctor, 2013a; Pfingstl and Kerschbaumer, 2022; Więcek et al., 2022). Mating behaviour has been studied in several groups, particularly spider mites (Tetranychidae) (Oku, 2014), predatory mites (Phytoseiidae) (Amano and Chant, 1979; Schulten, 1985; Ullah et al., 2017), and water mites (Mitchell, 1957; Więcek et al., 2022). However, detailed descriptions remain limited for many other groups, restricting comparisons of mating behaviour and our understanding of their reproductive biology.

The Acaridae comprise a diverse family of mites, many of which are fungivorous and occur in stored products and decaying organic matter (Fan and Zhang, 2007; Zhang et al., 2025b). Other species inhabit plants, insects and vertebrate nests, and some are economically important pests of stored products and agricultural crops (Díaz et al., 2000; Zhang et al., 2025b). Despite their ecological and economic importance, the mating behaviour of many acarid species remains insufficiently characterised. In Acaridae the male and female typically adopt opposing orientations during copulation, with the male’s ventral surface facing the female’s dorsum (Walter and Proctor, 2013a). Males also possess specialised structures, including anal suckers and paired suckers on tarsus IV, which help maintain contact with the female (Fan and Zhang, 2007; Walter and Proctor, 2013a). The male aedeagus is situated on the ventral idiosoma and is involved in direct insemination (Fan and Zhang, 2007; Witaliński, 2014). Describing the sequence of movements leading to copulation would provide a basis for understanding how these structures function during mating and for comparing mating behaviour within Acaridae and across mite groups.

*Tyrophagus curvipenis* Fain & Fauvel (Acari: Acaridae) has been recorded from various plant and animal hosts in Australia, France, Portugal, and New Zealand (Fan and Zhang, 2007; Ye and Zhang, 2014). This species was considered a better alternative diet than major pests such as *T. putrescentiae* for rearing predatory mites because it is naturally plant-inhabiting and thus may sustain predatory mite populations when the target pest is absent (Xie et al., 2018; Li et al., 2021; Xie et al., 2025). Its life cycle comprises five stages: egg, larva, protonymph, tritonymph, and adult. Under favourable conditions development from egg to adult takes approximately 10 days, and adults live for about 1 month and produce around 100 eggs during their lifetime (Ye and Zhang, 2014; Li et al., 2024). Diet also influences development and adult body size in this species (Ye and Zhang, 2014). Despite this basic knowledge of its life history, the mating behaviour and mate choice of *T. curvipenis* remain largely uncharacterised. Previous research has shown that mating access and social-sexual exposure are associated with female survival and reproduction, with unrestricted mating opportunities associated with higher lifetime fecundity but shorter lifespan than a treatment in which mating was limited to 24 h (Li et al., 2024). However, the number and timing of mating events were not controlled, leaving the effects of mating frequency on female reproductive performance unresolved.

Mate choice may depend on both the relatedness and previous mating history of potential partners. Kin discrimination can contribute to inbreeding avoidance when mating between relatives reduces offspring fitness (Hamilton, 1964; Mateo, 2004; Penn and Frommen, 2010). However, evidence of kin discrimination varies among mite species. For example, female *Tetranychus urticae* have been reported to prefer unrelated males over brothers (Tien et al., 2011), whereas no significant preference for siblings or unrelated partners was found in *Phytoseiulus persimilis* (Enigl and Schausberger, 2004). Whether *T. curvipenis* discriminates between kin and non-kin remains unknown. Previous mating history may also influence mate choice, because virgin and previously mated individuals may differ in their attractiveness or willingness to mate (Tsunoda and Amano, 2001; Balfour et al., 2026). Examining responses to relatedness and mating status would establish whether these factors are associated with mate choice in *T. curvipenis*.

The number of mates available to a female may also influence reproductive output. Polyandry, in which a female mates with multiple males, can increase fecundity through additional sperm transfer or other effects of mating, but repeated mating can also impose costs, including reduced survival of both sexes (Chapman et al., 1998; Arnqvist and Nilsson, 2000; Härdling and Kaitala, 2005; Li and Zhang, 2021). Multiple mating is common among acarid mites, although the reproductive consequences may depend on the frequency and circumstances of mating (Radwan and Rysińska, 1999). For example, in the acarids *Caloglyphus* (= *Sancassania*) *berlesei* and *Rhizoglyphus robini*, excessive mating can reduce female fitness (Radwan and Rysińska, 1999; Kołodziejczyk and Radwan, 2003). Additional mating has been associated with increased fecundity and a longer oviposition period in the predatory mites *Neoseiulus cucumeris* and *Amblyseius* (= *Neoseiulus*) *womersleyi* (Tsunoda and Amano, 2001; Ji et al., 2007). In *T. curvipenis*, Li et al. (2024) showed that mating access was associated with increased lifetime fecundity and reduced lifespan. However, their study did not establish how a controlled number of mating events affects female reproductive output. Comparing females that mate once, twice at different times in life, or have unlimited mating opportunities allows the effects of a second mating to be distinguished from those associated with unrestricted access to potential mates.

In this study we investigated mating behaviour, mate choice, and female reproductive performance in *T. curvipenis* through three complementary experiments. First, we described the mating sequence and developed an ethogram of mating behaviour. Second, we examined mate choice in relation to kinship by comparing responses to siblings and unrelated individuals. Third, we investigated mate choice in relation to partner mating status and assessed the effects of mating frequency on female reproductive performance. We compared responses to virgin and previously mated partners and measured female pre-oviposition period, oviposition period, daily oviposition rate, and fecundity under three mating treatments: one mating, two matings, and unlimited mating opportunities. Together, these experiments characterise mating behaviour and mate choice in *T. curvipenis* and examine how mating opportunities are associated with female reproductive output.

## 2 Materials and Methods

### 2.1 Mite culture

The stock culture of *T. curvipenis* was established from mites collected from capsicum (*Capsicum* sp.) leaves in a greenhouse at the Bioeconomy Science Institute Maiangi Taiao, St Johns, Auckland, New Zealand. The culture had been maintained in the laboratory for more than 2 years before the experiments. Mites were reared on abundant dry yeast pellets (*Saccharomyces cerevisiae*; Goodman Fielder Limited, New Zealand), which were sprinkled onto a black plastic sheet approximately 120 mm in diameter (Figure 1A). The sheet was placed on several layers of water-saturated tissue paper in a Petri dish (150 mm in diameter). Water was added as required to maintain moisture and high humidity and to prevent mites from escaping. The culture was maintained at 25 ± 1 °C, 80 ± 5% relative humidity and a photoperiod of 16:8 h (light:dark).

**Figure 1.**
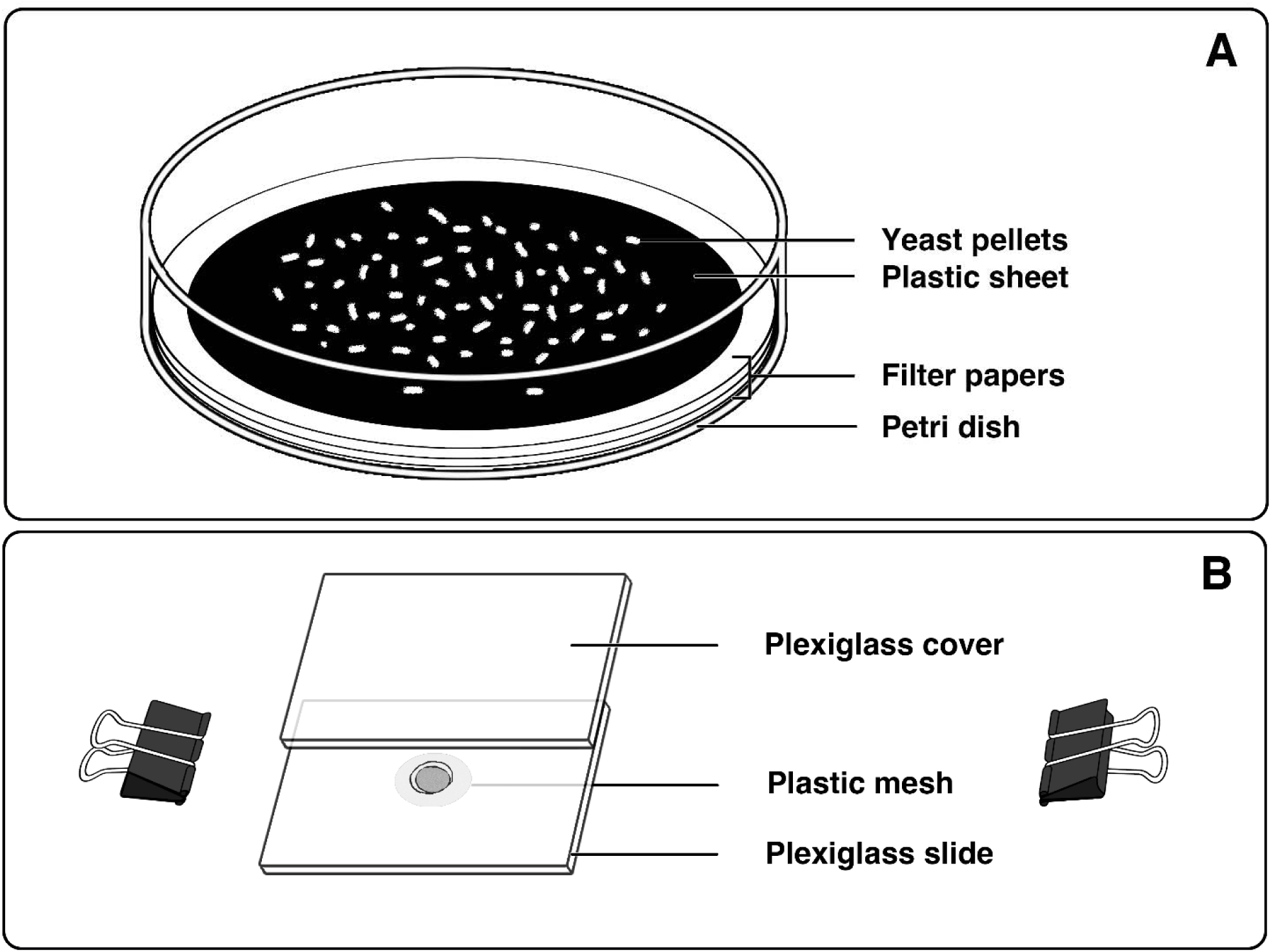
Experimental set-up. (A) Rearing set-up for the stock culture. (B) Modified Munger cell used in experiments. Illustrations were drawn by K. Zhang using Fresco and edited in Photoshop (Adobe, USA).

### 2.2 Experimental preparation

Modified Munger cells were used in all experiments (Wang et al., 2024). The cells consisted of two plexiglass slides (38 mm × 25 mm × 2 mm; length × width × thickness; Figure 1B). The lower slide contained a centrally located, truncated cone-shaped hole that served as the mite arena (top diameter = 5 mm; bottom diameter = 3 mm; volume = 25.66 mm³). The underside of the hole was covered with white plastic mesh secured with non-toxic PVC glue. The two slides were held together with metal clips.

Experimental mites were obtained through individual rearing. Gravid females were randomly selected from the stock culture and transferred individually to modified Munger cells, where abundant yeast pellets were provided as food. Eggs were collected daily and transferred individually to new cells. Individuals were monitored daily throughout development, and their sex was determined at adulthood using morphological characteristics under a dissecting microscope. Males were distinguished from females by their smaller body size and the presence of paired anal suckers on the posterior end of the venter (Fan and Zhang, 2007).

All adults used in the experiments were 3 days post-emergence. This age was selected to standardise adult age while allowing individuals to reach sexual maturity. For the kin-discrimination experiments (Part 2), related individuals were obtained from the same family, whereas unrelated individuals were obtained from different cultures (maintained separately). Eggs from the same family were reared individually to adulthood to establish their relatedness. The ovipositing females used to establish these families were allowed to lay eggs for no more than 1 week, with the males and females kept together during this period.

All individuals used in Parts 1 and 2 were virgin. Individuals used in Parts 1 and 3 were unrelated to their experimental partners. Both virgin and non-virgin individuals were used in Part 3. All experiments were conducted under the same temperature, relative humidity, and photoperiod conditions as the stock culture.

### 2.3 Part 1: Mating behaviour

A virgin female and a virgin male were introduced into a modified Munger cell and observed continuously under a dissecting microscope. Observations began immediately after introduction and continued until copulation commenced or for a maximum of 30 min. Pairs that did not mate within this period were excluded from analyses of mating behaviour and mating duration.

The mating sequence was recorded, and pre-copulation latency and copulation duration were measured to the nearest minute. Pre-copulation latency was defined as the time from the introduction of the male and female into the cell until the start of copulation. Copulation duration was defined as the time from the establishment of the copulatory position until the male and female separated. Female acceptance was characterised by the female remaining relatively stationary or moving slowly after the male mounted her, without actively attempting to escape.

A total of 95 pairs were observed, of which 89 mated within 30 min. For pairs that mated within this period, mating status was checked at 2 min intervals throughout copulation. No food was provided during mating.

### 2.4 Part 2: Kin discrimination in mate choice

Each trial involved one focal mite and two potential mates of the opposite sex: one sibling and one unrelated individual. The two potential mates were placed in a modified Munger cell before the focal mite was introduced. The mites were observed continuously under a dissecting microscope for a maximum of 40 min. The selected partner, pre-copulation latency, and copulation duration were recorded. The behaviour of the non-selected individual was also monitored, particularly attempts to interrupt copulation. Trials in which no mate choice occurred within 40 min were excluded from the mate-choice analysis. No food was provided during the choice test.

Two types of mate-choice trials were conducted:

1. **Male choice between sibling and unrelated females.** One potential mate was the focal male’s sister and the other was an unrelated female. Of 42 trials, 40 resulted in a mate choice within 40 min.
2. **Female choice between sibling and unrelated males.** One potential mate was the focal female’s brother and the other was an unrelated male. Of 40 trials, 38 resulted in a mate choice within 40 min.

To distinguish individuals within each trial, mites were marked with small watercolour dots on the dorsal surface before the experiment. Colours were assigned randomly to avoid confounding individual identity with marking colour.

### 2.5 Part 3: Mating status and reproduction

#### 2.5.1 Mate choice between virgin and previously mated partners

The experimental procedure followed that described in section 2.4, except that each focal mite was presented with one virgin and one previously mated potential partner of the opposite sex. Previously mated individuals were obtained from Part 1 and placed individually in modified Munger cells after their first mating. No food was provided during the choice test.

Two types of mate-choice trials were conducted:

1. **Male choice between virgin and previously mated females.** The previously mated female had mated once, 1 day before the trial, and was 4 days post-emergence at the time of testing. Of 40 trials, 35 resulted in a mate choice within 40 min.
2. **Female choice between virgin and previously mated males.** The previously mated male had mated once, 1 day before the trial, and was 4 days post-emergence at the time of testing. Of 42 trials, 40 resulted in a mate choice within 40 min.

#### 2.5.2 Mating frequency and reproductive performance

To investigate the effects of mating frequency on female reproductive performance, females were randomly assigned to one of three mating treatments, with 40 females per treatment. Abundant yeast pellets were provided and replenished when required.

1. **Single mating.** One virgin female and one virgin male were introduced into a modified Munger cell. The male was removed immediately after the first copulation ended.
2. **Two matings.** The procedure was the same as for the single-mating treatment, except that a second virgin male was introduced 6 days after the first mating. The second male was removed immediately after the second copulation ended.
3. **Unlimited mating opportunities.** One virgin female and one virgin male were kept together throughout the female’s reproductive period. The pair was not separated after copulation, allowing repeated mating. The number of observed mating events was recorded.

Females were checked at 24 h intervals, and the number of eggs laid each day was recorded and removed until oviposition ceased. For each female the pre-oviposition period, oviposition period, daily oviposition rate, and total fecundity were calculated. For females in the single-mating treatment, copulation duration was also recorded to examine its association with oviposition period and total fecundity. In the unlimited-opportunity treatment, females were observed mating an average of four times, with a range of two to eight observed mating events.

### 2.6 Statistical analysis

All analyses were conducted in R (R Core Team, 2026) in RStudio (version 2026.09.0). Data visualisation was performed using *ggplot2* (Wickham, 2016). Data were summarised as mean ± standard error of the mean (SE).

For Parts 2 and 3, mate-choice preferences were analysed using generalised linear models (GLMs) with a binomial distribution, testing whether the probability of choosing one partner type differed from 0.5. Pre-copulation latency and copulation duration were analysed using linear models (LMs), with focal sex and partner relatedness (Part 2) or partner mating status (Part 3) as fixed factors. The interaction between focal sex and partner type was also evaluated, where appropriate.

For the mating-frequency experiment (Part 3), pre-oviposition period was analysed using a GLM with a gamma distribution because the data were right-skewed. Oviposition period, daily oviposition rate and total fecundity were analysed using LMs, with mating treatment as the fixed factor. In the single-mating treatment, the relationships between copulation duration and oviposition period and between copulation duration and total fecundity were examined separately.

Model terms were evaluated using the *car* package (Fox and Weisberg, 2019). Where appropriate, significant effects were followed by estimated marginal means and pairwise comparisons with Holm adjustment for multiple comparisons using *emmeans* (Lenth, 2025). Data distributions were initially checked using histograms to determine model use. Model assumptions were then assessed using residual diagnostics, and dispersions were confirmed where applicable. All statistical tests used a significance threshold of α = 0.05.

## 3 Results

### 3.1 Part 1: Mating behaviour

The mating behaviour of *T. curvipenis* was divided into five steps: (I) encounter and recognition, (II) dorsal mounting, (III) attaining the copulatory position, (IV) copulation and sperm transfer, and (V) separation (Figure 2). Among successful matings, mean pre-copulation latency was 16.1 ± 0.9 min, and mean copulation duration (step IV) was 141.2 ± 3.0 min.

**Figure 2.**
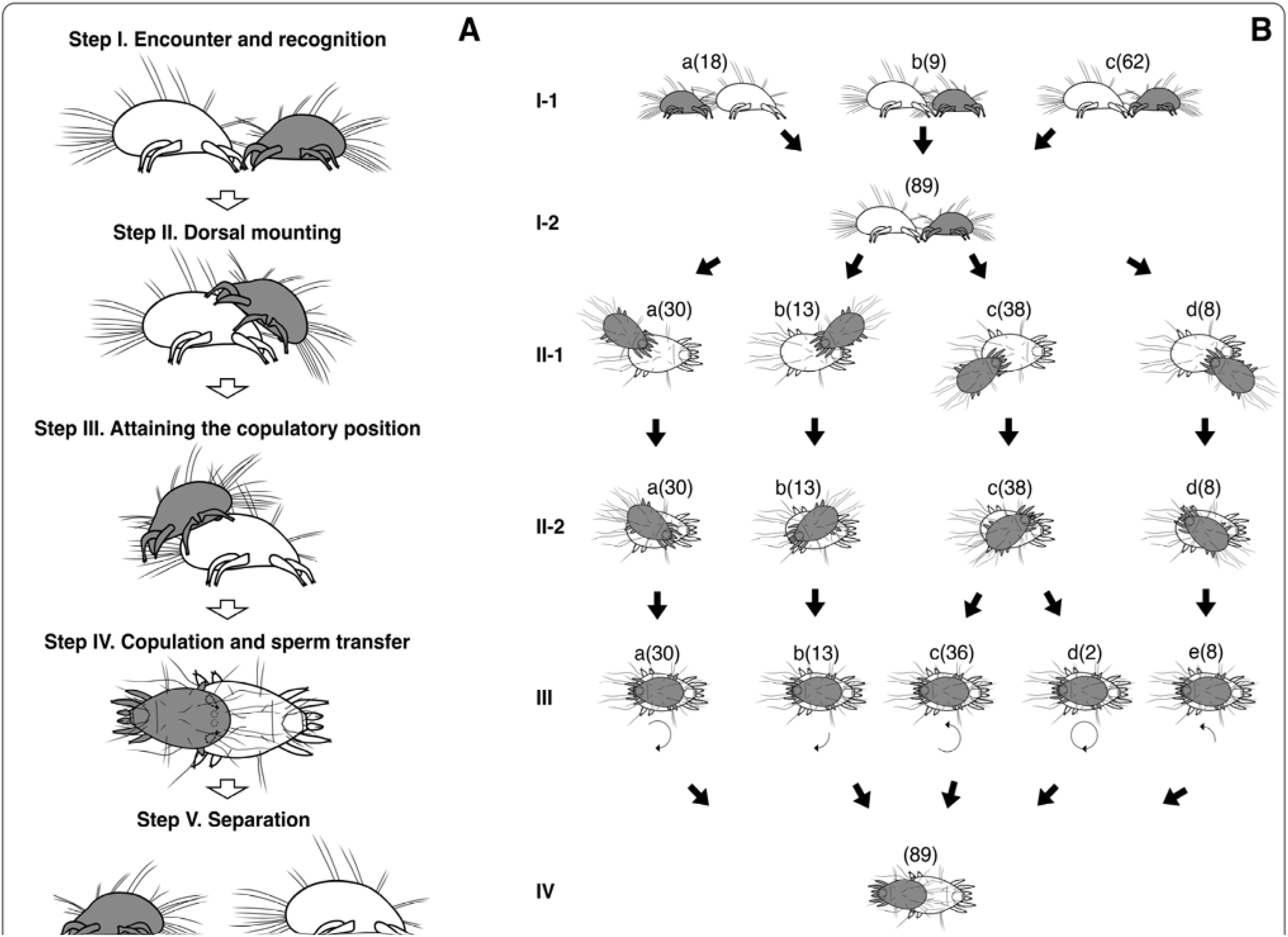
Ethogram of the mating behaviour of *Tyrophagus curvipenis*. (A) Schematic representation of the five steps of mating behaviour. In step IV, dashed lines indicate structures located beneath the male’s body, including a pair of leg IV and anal suckers. (B) Observed encounter types, mounting positions, and alternative movement patterns within each step. Numbers in parentheses indicate the number of observations for each pattern. In Step III, curved arrows indicate the direction and relative extent of male turning. Females are shown in white and males in grey. Illustrations were drawn by K. Zhang using Fresco and edited in Photoshop (Adobe, USA).

#### Step I: Encounter and recognition

Three types of initial encounter were observed: the male contacted the posterior end of a moving or stationary female (Figure 2B: I-1a; 20.2%); the female contacted the posterior end of a moving or stationary male (Figure 2B: I-1b; 10.1%); or the anterior ends of the male and female came into contact while one or both were moving (Figure 2B: I-1c; 69.7%). After initial contact, pairs often made repeated contact using their forelegs and palps (Figure 2A, B: I-2). Females sometimes moved away rapidly or vigorously after brief contact. In some cases, males departed while females remained relatively stationary. When both individuals remained in place, the male generally proceeded to mount the female.

#### Step II: Dorsal mounting

Males mounted females from four positions: left posterior (Figure 2B: II-1a; 33.7%), left anterior (Figure 2B: II-1b; 14.6%), right posterior (Figure 2B: II-1c; 42.7%), and right anterior (Figure 2B: II-1d; 9.0%). Females generally remained stationary while males climbed onto their dorsum. Occasionally, females continued moving vigorously, preventing successful mounting or causing males to fall off after mounting.

#### Step III: Attaining the copulatory position

After mounting the female’s dorsum, the male repositioned itself into a tail-to-tail arrangement, with the male and female facing in opposite directions. The direction and extent of turning depended on the male’s initial position on the female. Males generally appeared to take the shorter route, involving a smaller turning angle, to reach the copulatory position. However, two males turned through an angle greater than 180° (Figure 2B: III-d; 2.2%).

#### Step IV: Copulation and sperm transfer

During copulation, males maintained contact with females using paired suckers (Figure 2A: IV). In this position, the male’s aedeagus, located near the centre of the ventral idiosoma, was close to the female’s copulatory opening near the posterior end of the idiosoma. However, the movements of the aedeagus and the process of insemination could not be observed because of the mating position and the limited magnification of the dissecting microscope. The fourth pair of legs (leg IV) was usually curled inward and not visible in dorsal view. These legs showed little movement apart from occasional tapping. The third pair of legs (leg III) repeatedly tapped the female’s body, whereas the first and second pairs (legs I and II) mainly supported the male and helped maintain its position by contacting the substrate. Females generally remained stationary during copulation.

#### Step V: Separation

At the end of copulation the male and female separated and departed without further contact.

### 3.2 Part 2: Kin discrimination in mate choice

Neither males nor females showed a significant preference for siblings or unrelated individuals as mating partners (GLM: *z* = −0.32, *P* = 0.752 for males; *z* = −0.32, *P* = 0.746 for females; Figure 3). Mating preferences did not differ significantly between the sexes (Wald χ^2^ < 0.001, *df* = 1, *P* = 0.991).

**Figure 3.**
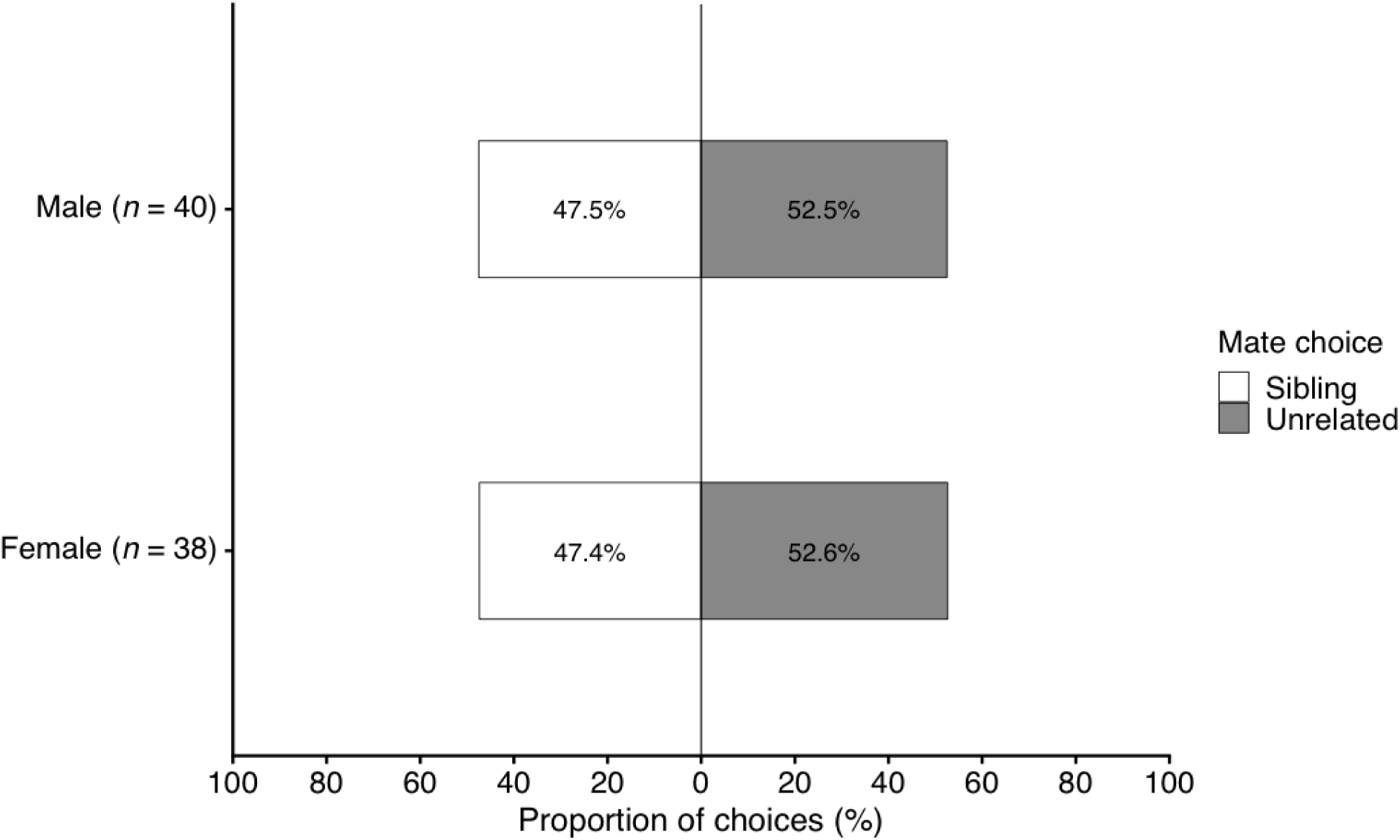
Proportion of mating choices by male and female *Tyrophagus curvipenis* when each focal individual was given a choice between a sibling and an unrelated individual of the opposite sex.

Pre-copulation latency (LM: *F*_1,74_ = 0.19, *P* = 0.892) and copulation duration (*F*_1,74_ = 0.13, *P* = 0.717) were not significantly affected by relatedness of the mating partner. Both durations, however, were significantly affected by the sex of the focal subject (pre-copulation latency: *F*_1,74_ = 16.00, *P* < 0.001; copulation duration: *F*_1,74_ = 10.01, *P* = 0.002). Pre-copulation latency was shorter when males were the focal subjects than when females were the focal subjects (Table 1). Although copulation duration was also shorter for males, the pairwise comparisons did not detect significant differences among the four groups. No significant interaction was detected between focal subject sex and partner relatedness for either pre-copulation latency (*F*_1,74_ = 0.31, *P* = 0.577) or copulation duration (*F*_1,74_ < 0.001, *P* = 0.997).

**Table 1.** Pre-copulation latency and copulation duration (min) of *Tyrophagus curvipenis* in mating trials in which each focal subject was given a choice between a sibling and an unrelated individual of the opposite sex. Values are means ± SE. Within each response variable, means sharing no letters differ significantly based on estimated marginal means and Holm-adjusted pairwise comparisons (*P* < 0.05).

| Focal subject | Mate relatedness | *n* | Pre-copulation | Copulation |
|---|---|---|---|---|
| Male | Sibling | 19 | 20.1 ± 1.5<sup>ab</sup> | 136.8 ± 6.4<sup>a</sup> |
|  | Unrelated | 21 | 19.1 ± 1.4<sup>a</sup> | 138.6 ± 4.7<sup>a</sup> |
| Female | Sibling | 18 | 24.8 ± 1.4<sup>bc</sup> | 121.6 ± 3.8<sup>a</sup> |
|  | Unrelated | 20 | 25.5 ± 1.3<sup>c</sup> | 123.4 ± 4.0<sup>a</sup> |

### 3.3 Part 3: Mating status and polyandry

#### 3.3.1 Mate choice for virgin or mated partners

Males and females differed in their mating preferences when given a choice between a virgin and a previously mated partner (Figure 4). Males showed a significant preference for virgin partners (*z* = 3.85, *P* < 0.001), whereas females showed no significant preferences for either partner (*z* = 1.87, *P* = 0.062).

**Figure 4.**
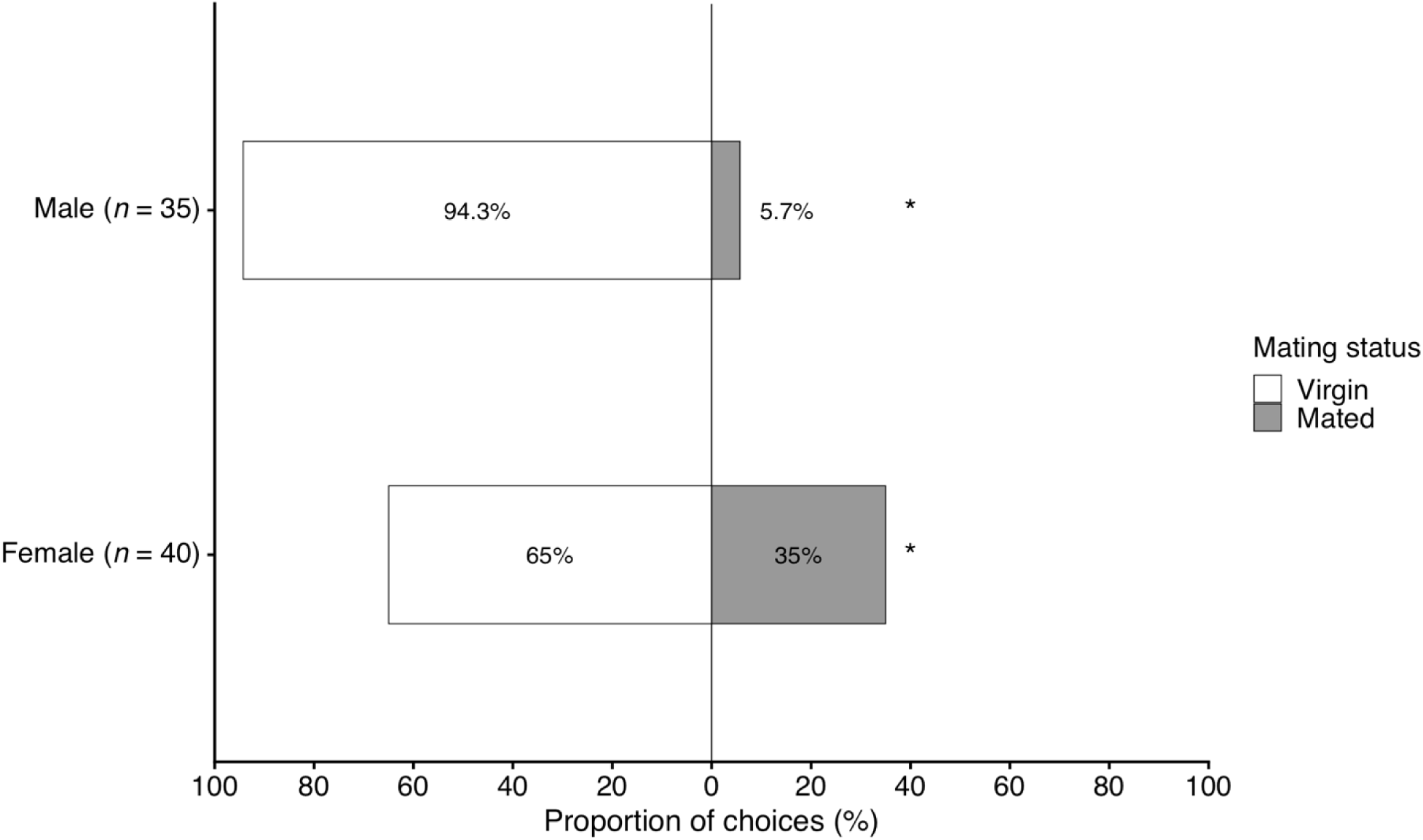
Proportion of mating choices by male and female *Tyrophagus curvipenis* when each focal individual was given a choice between a virgin and a previously mated individual of the opposite sex. Asterisks indicate a significant difference between the sexes in mating preference (Wald χ^2^ = 10.62, *df* = 1, *P* = 0.001).

Pre-copulation latency was significantly affected by focal subject sex (LM: *F*_1,71_ = 14.95, *P* < 0.001) and the interaction between focal subject sex and partner mating status (*F*_1,71_ = 10.98, *P* = 0.001), but not by partner mating status alone (*F*_1,71_ = 1.84, *P* = 0.180). Pre-copulation latency was shorter when males were the focal subjects, and the effect of partner mating status differed between the sexes (Table 2).

**Table 2.** Pre-copulation latency and copulation duration (min) of *Tyrophagus curvipenis* in mating trials in which each focal subject was given a choice between a virgin and a previously mated individual of the opposite sex. Values are means ± SE. Within each response variable, means sharing no letters differ significantly based on estimated marginal means and Holm-adjusted pairwise comparisons (*P* < 0.05).

| Focal subject | Mating status of partner | *n* | Pre-copulation | Copulation |
|---|---|---|---|---|
| Male | Virgin | 33 | 17.3 ± 1.0<sup>a</sup> | 140.6 ± 3.9<sup>c</sup> |
|  | Mated | 2 | 27.0 ± 1.0<sup>ab</sup> | 102.0 ± 10.0<sup>ab</sup> |
| Female | Virgin | 26 | 24.3 ± 1.1<sup>b</sup> | 127.3 ± 3.9<sup>b</sup> |
|  | Mated | 14 | 19.6 ± 1.1<sup>ab</sup> | 106.6 ± 4.0<sup>a</sup> |

Copulation duration was significantly affected by focal subject sex (*F*_1,71_ = 15.48, *P* < 0.001) and partner mating status (*F*_1,71_ = 15.21, *P* < 0.001), but their interaction was not significant (*F*_1,71_ = 1.24, *P* = 0.269). Copulation duration was shorter when females were the focal subjects and when the partners were mated (Table 2).

#### 3.3.2 Mating frequency and reproductive performance

Pre-oviposition period was not significantly affected by mating frequency (GLM: Wald χ^2^ = 1.97, *df* = 2, *P* = 0.373; Table 3). Oviposition period increased significantly with mating frequency (LM: *F*_2,117_ = 38.96, *P* < 0.001; Table 3). Daily oviposition also differed significantly among treatments (*F*_2,117_ = 5.65, *P* = 0.005), with females mated twice laying fewer eggs per day than females given a single mating or unlimited mating opportunities (Table 3). Fecundity increased significantly with mating frequency (*F*_2,117_ = 42.02, *P* < 0.001; Table 3). Among females that mated once, copulation duration was positively associated with both oviposition period and total fecundity (Figure 5).

**Figure 5.**
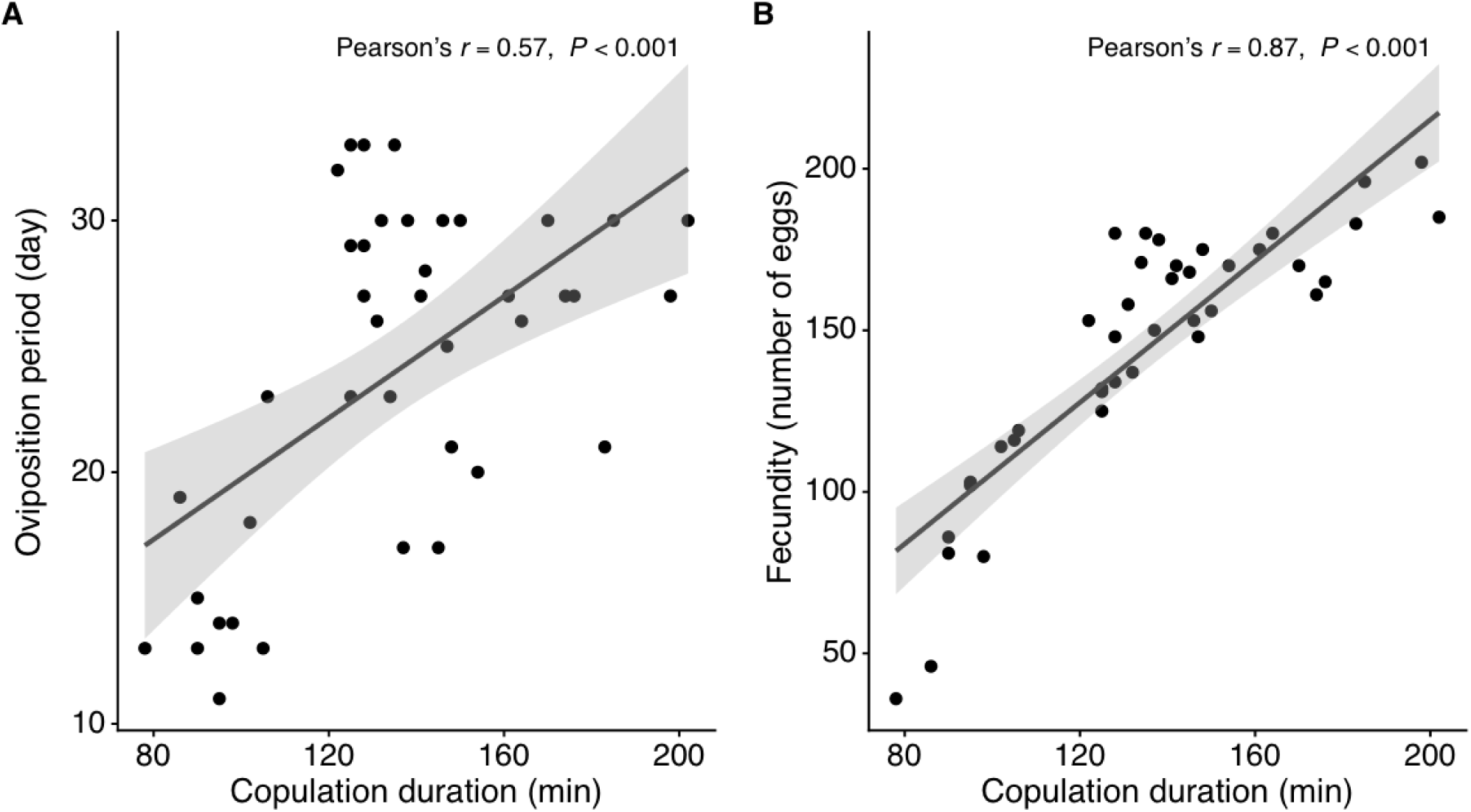
Relationships between copulation duration and reproductive performance in female *Tyrophagus curvipenis* that mated once. (A) Oviposition period. (B) Total fecundity. Pearson’s correlation coefficients and corresponding *P*-values are shown in each panel.

**Table 3.** Reproductive parameters of female *Tyrophagus curvipenis* under three mating-frequency treatments: a single mating, two matings, and unlimited mating opportunities. Values are means ± SE. Within each response variable, means sharing no letters differ significantly based on estimated marginal means and Holm-adjusted pairwise comparisons (*P* < 0.05).

| Mating frequency | *n* | Pre-oviposition (days) | Oviposition (days) | Daily oviposition (eggs/day) | Fecundity (number of eggs) |
|---|---|---|---|---|---|
| Single mate | 40 | 1.2 ± 0.1<sup>a</sup> | 24.0 ± 1.0<sup>a</sup> | 6.2 ± 0.3<sup>b</sup> | 144.6 ± 6.2<sup>a</sup> |
| Two mates | 40 | 1.2 ± 0.1<sup>a</sup> | 32.1 ± 1.1<sup>b</sup> | 5.3 ± 0.1<sup>a</sup> | 168.5 ± 6.2<sup>b</sup> |
| Unlimited mating | 40 | 1.3 ± 0.1<sup>a</sup> | 39.8 ± 1.6<sup>c</sup> | 6.0 ± 0.2<sup>b</sup> | 235.0 ± 8.9<sup>c</sup> |

## 4 Discussion

This study describes the mating behaviour of *T. curvipenis* and examines mate choice in relation to kinship and mating status, as well as the effects of mating treatment on female reproductive performance. Mating followed a sequence of initial contact, dorsal mounting, repositioning, and copulation. Neither sex showed a significant preference for siblings or unrelated partners, whereas males preferred virgin over previously mated females. Mating treatment affected oviposition period, daily oviposition rate, and total fecundity, with females given unlimited mating opportunities producing the most eggs. These findings provide a basis for further investigation of mating behaviour and reproductive performance in this species.

### 4.1 Mating behaviour

The mating sequence of *T. curvipenis* involved initial contact, dorsal mounting, and repositioning into a tail-to-tail copulatory position, which is consistent with other reported species within this family (Levinson et al., 1989; Walter and Proctor, 2013a). Males of *T. curvipenis* mounted females from different positions, indicating that mating could proceed from several initial orientations. Initial encounters most often involved contact between the anterior ends, frequently followed by repeated contact using the forelegs and palps. Although these observations characterise the sequence of mating behaviour, the cues involved in partner recognition and acceptance were not determined. Chemical cues, including sex pheromones, have been implicated in mating behaviour in other acarid mites (Sonenshine, 1985; Levinson et al., 1989; Mori et al., 1998; Ryono et al., 2001; Mizoguchi et al., 2003), but their role in *T. curvipenis* remains unknown.

Mean copulation duration in *T. curvipenis* was approximately 140 min, which is considerably longer than the pre-copulation period. This duration is comparable to the 128–219 min reported for *Acarus siro* (Levinson et al., 1989). Copulation can also last several hours in some phytoseiid and water mites (Proctor and Smith, 1994; Momen, 1997; Więcek et al., 2022; Zhang et al., 2025a), whereas considerably shorter durations have been reported in tetranychid and tenuipalpid mites (Aponte and McMurtry, 1997; Gomes et al., 2026). The reasons for these differences remain unclear. Whether copulation duration reflects differences in insemination mode, sperm storage systems and lifetime sperm requirements, lifestyle, or predation risk should be investigated further.

### 4.2 Kin discrimination and mating interactions

In this study neither males nor females showed a significant preference for siblings or unrelated partners, and partner relatedness did not affect pre-copulation latency or copulation duration. These results provide no evidence of kin discrimination under the conditions tested. The response was similar to that found for *P. persimilis*, which showed no significant preference for siblings or unrelated partners (Enigl and Schausberger, 2004). Empirical evidence suggests that kin discrimination is not a static trait in mite mating systems, and inbreeding avoidance can be deprioritised relative to immediate reproductive opportunities (Schausberger and Sato, 2020; Zhou et al., 2020). Nevertheless, the present results do not establish whether this species can recognise kin under other conditions.

However, kin recognition can depend on learning and familiarity rather than innate recognition alone (Penn and Frommen, 2010). Because individuals in this experiment were reared separately, *T. curvipenis* may have had limited opportunities to become familiar with their kin, potentially reducing their ability to discriminate between siblings and unrelated individuals under more natural conditions (Schausberger, 2007; Zhang and Zhang, 2022). Thus, the absence of a significant preference for either group does not necessarily exclude kin discrimination under other rearing conditions and the absence of kin recognition ability.

Competition between potential mates was also observed during the choice trials (S.-S. Ye, pers. obs.). Males frequently fought before contacting females and sometimes interfered with ongoing copulation, whereas female–female fighting was rare. These interactions may have influenced the observed mating outcomes, particularly in trials where females were presented with two males. Consequently, the partner that mated may not always have been the female’s preferred choice. This was also reflected by a longer pre-copulation latency and short copulation duration when females were the focal subject. Further experiments that separate mate preference from competition would help clarify their respective effects.

### 4.3 Mate choice in relation to mating status

In this study males significantly preferred virgin females, whereas females showed no significant preference between virgin and previously mated males. This result suggests that partner mating status was associated with male choice under the conditions tested, but not with a statistically significant preference by females. In spider mites sperm precedence can favour the first male to mate, potentially increasing the reproductive value of mating with virgin females (Potter et al., 1976; Potter and Wrensch, 1978; Takafuji, 1986; Walter and Proctor, 2013a). However, sperm precedence has not been established in *T. curvipenis* and other acarid mite species (Klimov and Sidorchuk, 2011), so its relevance to the observed male preference remains unknown.

During the trials, previously mated females were sometimes less active and remained apart while males interacted with virgin females (S.-S. Ye, pers. obs.). This observation raises the possibility that differences in female receptivity contributed to the observed preference. For example, females of *Amblyseius* (= *Neoseiulus*) *womersleyi* mated 1 day before tend to reject mating on the 2nd day (Tsunoda and Amano, 2001). Because male choice and female receptivity were not experimentally separated, the extent to which males actively discriminated between mating-status classes cannot be determined.

Copulation duration was shorter with previously mated partners, with no significant interaction between partner mating status and focal subject sex. A similar pattern has been reported in the spider mite *Amphitetranychus viennensis*, in which males copulated for less time with fertilised females than with virgins (Yang et al., 2021). However, the number of trials involving males and previously mated females in the present study was very small, limiting interpretation of this comparison. Moreover, shorter copulation does not necessarily indicate reduced sperm transfer or fertilisation success (Zhang et al., 2025a). Direct measurements would be needed to establish the reproductive consequences of mating with previously mated partners.

### 4.4 Mating treatment and female reproductive performance

Mating treatment affected oviposition period, daily oviposition, and fecundity, but not the pre-oviposition period. Females given unlimited mating opportunities had the longest oviposition period and highest fecundity. Females that mated twice had a lower daily oviposition than those that mated once or had unlimited mating opportunities. Thus, differences in fecundity were associated with differences in oviposition period rather than a consistent increase in daily egg production across treatments. As discussed earlier, similar reproductive patterns have been reported in other mites, where additional mating increased fecundity and oviposition period (Elbadry and Elbenhawy, 1968; Ji et al., 2007). Out results suggest that *T. curvipenis* females require multiple matings to reach maximum reproduction.

Mating twice resulted in a lower daily oviposition rate. This may be because the second mating extended the oviposition period without affecting egg production earlier in the period, resulting in a greater increase in oviposition duration than in the total number of eggs laid and, consequently, a lower daily oviposition rate.

Among females that mated once, longer copulation was associated with a longer oviposition period and greater total fecundity. Although longer copulation may be associated with greater sperm transfer in some mites (Schulten et al., 1978; Ullah et al., 2017), the present correlations do not establish this mechanism. Copulation duration may also reflect other differences between mating pairs. Measurements of sperm transfer and fertilisation would be needed to determine whether copulation duration predicts female reproductive output.

### 4.5 Conclusion

This study characterised the mating sequence of *T. curvipenis* and examined mate choice and female reproductive performance, providing a basis for further research on mating systems in mites. The absence of detectable kin discrimination under the experimental conditions raises the question of whether kin recognition occurs under more natural conditions, where individuals may have opportunities to become familiar with their relatives. The shorter copulation duration with previously mated females also merits further investigation to determine whether it reflects differences in female receptivity, sperm transfer or other aspects of mating. Furthermore, the higher total fecundity under unlimited mating opportunities suggests that repeated mating may contribute to female reproductive output. Future studies should also examine whether mating with different males affects female fecundity. These investigations would provide further insight into mate choice and the reproductive significance of polyandry in *T. curvipenis*.

## 6 Author Contributions

SY: Investigation, Methodology, Writing – original draft, review & editing; KZ: Formal Analysis, Visualization, Writing – original draft, Writing – review & editing; ZQZ: Supervision, Methodology, Writing – review & editing, Project administration. SY and KZ contributed equally to this work and share first authorship. All authors contributed to the article and approved the submitted version.

## 7 Acknowledgements

We thank Ray Prebble (Bioeconomy Science Institute Maiangi Taiao, New Zealand) for reviewing an early version of this manuscript and for comments. Shi-Sen Ye was supported by the Training Program of Fujian Excellent Talents in Universities (FETU), which is greatly appreciated.

## 8 Data Availability Statement

The datasets analysed for this study can be found in the DataStore: https://datastore.landcareresearch.co.nz/dataset/tyrophaguscurvipenis-mating

